# Large scale *in vivo* recording of sensory neuron activity with GCaMP6

**DOI:** 10.1101/166959

**Authors:** Kim I Chisholm, Nikita Khovanov, Douglas M Lopes, Federica La Russa, Stephen B McMahon

**Author notes:** To whom correspondence should be addressed Neurorestoration group Wolfson Wing Hodgkin Building King’s College London Guy’s Campus, London Bridge London SE1 1UL. Author contributions: Conceptualisation, S.B.M. and K.I.C.; Methodology, S.B.M. and K.I.C.; Investigation, K.I.C, N.K., D.M.L, F.L.R.; Writing – Original Draft, S.B.M. and K.I.C. Writing – Review & Editing, S.B.M., N.K., D.M.L., F.L.R Funding Acquisition, S.B.M., Supervision, S.B.M., Formal Analysis, K.I.C.

## Abstract

Greater emphasis on the study of intact cellular networks in their physiological environment has led to rapid advances in intravital imaging in the central nervous system, while the peripheral system remains largely unexplored. To assess large networks of sensory neurons we selectively label primary afferents with GCaMP6s and visualise their functional responses *in vivo* to peripheral stimulation. We show that we are able to monitor simultaneously the activity of hundreds of sensory neurons with sensitivity sufficient to detect, in most cases, single action potentials with a typical rise time of around 200 milliseconds, and an exponential decay with a time constant of approximately 700 milliseconds. Using this sensitive technique we are able to show that large scale recordings demonstrate the recently disputed polymodality of nociceptive primary afferents with between 40-80% of thermally sensitive DRG neurons responding also to noxious mechanical stimulation. We also specifically assess the small population of peripheral cold fibres and demonstrate significant sensitisation to cooling after a model of sterile and persistent inflammation, with significantly increased sensitivity already at decreases of 5°C when compared to uninflamed responses. This not only reveals interesting new insights into the (patho)physiology of the peripheral nervous system but also demonstrates the sensitivity of this imaging technique to physiological changes in primary afferents.

**Significance Statement:** Most of our functional understanding of the peripheral nervous system has come from single unit recordings. However, the acquisition of such data is labour-intensive and usually ‘low yield’. Moreover, some questions are best addressed by studying populations of neurons. To this end we report on a system that monitors activity in hundreds of single sensory neurons simultaneously, with sufficient sensitivity to detect in most cases single action potentials. We use this technique to characterise nociceptor properties and demonstrate polymodality in the majority of neurons and their sensitization under inflammatory conditions. We therefore believe this approach will be very useful for the studies of the somatosensory system in general and pain in particular.

## Introduction

Primary somatosensory neurons are functionally diverse, with anatomical, physiological and molecular techniques identifying about one or two dozen subtypes (Kandel et al., 2013; Usoskin et al., 2014). These neurons are critical for a large number of distinct sensations including, but not limited to, touch, pain, itch, proprioception and temperature. Our knowledge regarding the encoding properties of these neurons is largely derived from single unit recording studies of individual afferent fibres, work that is hampered by the low throughput and inherent stochasticity of such data acquisition.

The advent of genetically encoded calcium indicators has opened up the possibility for large scale optical assessment of the functional and morphological characteristics of entire neuronal networks with good spatial and temporal sensitivity. These techniques have been applied to a variety of central nervous system (CNS) structures, including sensory, motor and visual cortex and spinal cord (Stosiek et al., 2003; Dombeck et al., 2007; Flusberg et al., 2008; Tian et al., 2009; Johannssen and Helmchen, 2010; Ghosh et al., 2011; Chen et al., 2012, 2013; Zariwala et al., 2012; Sun et al., 2013; Dana et al., 2014; Sekiguchi et al., 2016) and, very recently, some peripheral networks (Barretto et al., 2014; Williams et al., 2016; Wu et al., 2015). In recent months a number of papers have describe the application of *in vivo* imaging also to the dorsal root ganglia (Emery et al., 2016; Kim et al., 2016; Smith-Edwards et al., 2016) but due to the novelty of the technique the peripheral nervous system still remains relatively unexplored.

Indeed, it is evident from this small string of papers, that the application of this technique to the peripheral nervous system is still an evolving field. For example initially slower version of the calcium indicator GCaMP as well as very slow image acquisition speeds may have hampered a more detailed analysis of the pathophysiology of the peripheral nervous system (Kim et al., 2016; Smith-Edwards et al., 2016) while small numbers of sampled cells could reduce the benefits inherent to this approach (Emery et al., 2016; Smith-Edwards et al., 2016). One such evolving claim suggested that the vast majority of primary afferent exhibit modality specificity (Emery et al., 2016) in contrast to frequently observed nociceptive polymodality seen using microneurography, electrophysiology and multiple lines of *in vitro* evidence. Here we re-examine this claim using large scale *in vivo* GCaMP imaging, confirming traditional views of widespread nociceptive polymodality.

In this study we present an enhanced method which provides for the first time the added benefits of being able to sample from hundreds of cells simultaneously, at acquisition frequencies above 1 Hz, using very low light levels with no detectable photo-bleaching and an increased resistance to sample movement. These features make this technique particularly useful to the study of large populations of heterogeneous cells responding variably across time. We were therefore able to efficiently sample the diverse array of DRG neurons in response to acute and persistent pain states. Examining a very frequently used pain model we report on an unexpected involvement of the peripheral nervous system only in the initial phase of the formalin induced pain response. Owing to the large sampling capacity we were additionally able to examine rare populations of cells: in a model of persistent pain we are able to demonstrate a novel and significant sensitisation of cold nociceptors – a small population of cells difficult to examine using traditional recording techniques.

## Methods

For most experiments we used adult C57BL/6 mice (Envirgo) and expressed GCaMP6s in sensory neurons via intrathecal AAV9 (see below). For comparison, we also studied responses in Cre-dependent C57BL/6 GCaMP6s mice, (Jackson Labs, Ai96 line, Jackson Labs. Stock No: 028866) crossed with Advillin-CreERT2 (courtesy of John Wood), and also in Snap25-2A-GCaMP6s-D (Jackson Labs. Stock No 025111). The animals weighed 20-35g at the time of experimentation. Both male and female mice were housed on a 12hr light/dark cycle with a maximum of 8 mice per cage, with food and water available *ad libitum.* All experiments were performed in accordance with the UK Home Office Animals (Scientific Procedures) Act (1986).

### Intrathecal administration of AAV9-GCaMP6s

C57BL/6 mice were anaesthetised with isoflurane (~2% in oxygen) and Carprieve (0.025 mg; Norbrook Laboratories) was administered subcutaneously for post-operative pain management. Mice were maintained at around 37°C using a homeothermic heating mat. An incision was made in the skin over the lumbar region and muscle was removed to expose the intervertebral membrane between T12 and T13 vertebrae. The region surrounding the lumbar enlargement provides easier access to the intervertebral area and ensures minimal damage. A small cut was made in the membrane and the underlying dura in order to insert a small catheter of 0.2 mm diameter (Braintree Scientific) in the caudal direction, through which 5 μl of AAV9.CAG.GCaMP6s.WPRE.SV40 (UPENN Vector Core, AV-1-PV2833, 1.1x1013 gc/ml) was infused into the intrathecal space at 1.2 μl/min. Due to the length of the inserted cannula the infusion was close to the L4 DRGs. The catheter was left in place for 2 minutes before slow withdrawal. The incision was closed and mice were allowed to recover for between 2 weeks to up to 4 months.

### Tamoxifen dosing

Tamoxifen (T5648; Sigma-Aldrich) was dissolved in 100% ethanol to a concentration of 97.5μg/μl and further dissolved to a working concentration of 7.8μg/μl in wheat germ oil. This was placed on a shaker at room temperature for 2–3 hr and subsequently stored at –20°C. To induce GCaMP expression, mice carrying the Advillin-CreERT2 allele received 75mg/kg tamoxifen intraperitoneally (IP) once daily for 3 days. Gene expression was allowed to occur for a minimum of 14 days before mice were imaged.

### *In vivo* imaging of sensory neuron activity using GCaMP responses

For *in vivo* imaging, mice were anaesthetised using urethane (12.5% w/v). An initial dose of 37.5 mg (in a volume of 0.3 ml) was given IP. Further doses were given at approximately 15-20 minute intervals, depending on hind limb and corneal reflex activity, until surgical depth was achieved. The core body temperature was maintained close to 37°C using a homeothermic heating mat with a rectal probe (FHC). A tracheal catheter was installed and the mice breathed spontaneously. Animals were hydrated with 0.5 ml sterile normal saline (0.9%) administered subcutaneously. An incision was made in the skin on the back and the muscle overlying the L3, L4 and L5 DRG was removed. The bone around the L4 DRG was carefully removed in a caudal-rostral direction and the underlying epineurium and dura mater over the DRG were washed and moistened with normal saline. The position of the mouse was varied between prone and lateral recumbent to orient the DRG in a more horizontal plane. The exposure was then stabilised at the neighbouring vertebrae using spinal clamps (Precision Systems and Instrumentation) attached to a custom made imaging stage. The exposed cord and DRG were covered with silicone elastomer (World Precision Instruments, Ltd) to avoid drying and to maintain a physiological environment. The mouse was then placed under the Eclipse Ni-E FN upright confocal/multiphoton microscope (Nikon) and the microscope stage was variably diagonally orientated to optimise focus on the DRG. The ambient temperature during imaging was kept at 32°C throughout. All images were acquired using a 10X dry objective. To obtain confocal images a 488 nm Argon ion laser line was used, while a Coherent Chameleon II laser was tuned to 920 nm for multiphoton imaging. GCaMP signal was collected at 500-550 nm. Time series recordings were taken with an in-plane resolution of 512 × 512 pixels and a fully open pinhole for confocal image acquisition. Temporal image acquisition varied between 1-16 Hz depending on the experimental requirements and signal strength.

### Activation of sensory neurons with electrical stimuli

In some animals, the sciatic nerve on the side ipsilateral to the DRG being imaged was exposed through blunt dissection. A custom made cuff electrode with Teflon insulated silver wire (Ø 0.125 mm; Advent Research Materials) was placed underneath and then around the sciatic nerve. The preparation was isolated and stabilised using dental silicon impression compound (Heraeus Kulzer). A biphasic stimulator (World Precision Instruments) was used to deliver individual or trains of square wave current pulses to the sciatic nerve. Pulses of 250 μsec duration and 250 μA amplitude were used to activate myelinated (A) fibres in the sciatic nerve, and supramaximal pulses of 1 msec duration and 5 mA amplitude were used to activate all afferent fibres, both myelinated (A) and unmyelinated (C) axons.

### Activation of sensory neurons with thermal stimuli

A Peltier device (TSAII, Medoc) with a 16x16 mm probe was placed onto the plantar surface of the hind paw ipsilateral to the DRG being imaged. The temperature of the block was increased from a baseline temperature of 32°C to 50°C or decreased from 32°C to 4°C. Temperature changes occurred as either a ramp of 1.5°C/sec to the target temperature (maintained for 10 seconds) before returning back to baseline at 4°C/sec, or as increasing/decreasing incremental temperature ramps. Consecutive increments occurred as steps of 2°C in the case of an increase and 5°C during a decrease (with a final drop of 3°C from 7° to 4°C) and a minimum of 90 seconds between incremental changes. Each individual increment involved a temperature change of 2°C/sec, a holding temperature for 5 seconds and a return to baseline at 4°C/sec.

### Activation of sensory neurons with mechanical and chemical stimuli

Mechanical stimulation consisted of brushing or pinching of the plantar surface of the ipsilateral hind paw. To achieve the stimulation of the maximal number of receptive fields the entire plantar surface was pinched with blunt forceps across most of the surface of the paw. An effort was made to stimulate similar areas across different experiments in order to activate a comparable number of sensory fibres.

In some experiments, sterile normal saline or formalin (1.85% in saline) was injected at a volume of 20 μl into the middle of the plantar surface of the ipsilateral hind paw. Responses of L4 DRG neurons were monitored for at least 30 minutes after injection of saline and formalin.

### Behaviour assessment after formalin injection

For the behaviour experiments, mice were placed in a Perspex chamber on a wire mesh floor and allowed to acclimatise for at least 30 minutes. After acclimatisation, mice were lightly restrained and 20μl of 1.85% formalin in 0.9% saline solution was injected subcutaneously into the plantar surface of the hind paw using a 31 gauge syringe. The animal was then put back into the chamber and its behaviour was recorded for 30 minutes. Pain behaviour (time spent flinching, jerking or licking the injected paw) was quantified from video footage, using Etholog Software (Ottoni, 2000) in blocks of 5 minutes. All behavioural experiments were carried out during the light cycle of the day.

### UVB irradiation

Mice were anaesthetised with ketamine (75mg/kg in saline, Narketan, Vetoquinol) and medetomidine (0.5mg/kg in saline, Dormitor, Vetoquinol) and placed under a black cloth for protection. Their eyes were moistened and protected with eye gel (Viscotears, Liquid Gel, Novartis) and the plantar surface of the left paw was exposed through slits in the material and immobilised using tape. The paws of mice were exposed to 3000 mJ/cm^2^ over approximately 40 minutes (the duration of exposure depended on the strength of the UVB lamp -TL01 fluorescent bulbs (maximum wavelength 311nm)-, assessed before each irradiation using a photometer). Control animals were anaesthetised but not irradiated. Animals were allowed to recover for 48 hrs before *in vivo* imaging.

### Evaluation of transfection efficiency of DRGs

Fourteen days after intrathecal injection of AAV9-GCaMP6s, four mice were terminally anaesthetised and transcardially perfused with PBS followed by 4% paraformaldehyde (PFA). The L4 DRGs were removed and post-fixed for 2 hrs in 4% PFA before cryoprotection in 30% sucrose (with 0.02% sodium azide) for 24 hours. They were then embedded in Optimal Cutting Temperature (Tissue-Tek), cut into 10 μm sections at -20°C and mounted onto glass slides.

Once dried, DRG slices were rehydrated and blocked with 10% serum for one hour prior to incubation with primary antibodies against β-III-tubulin (primary afferent marker, 1:1000, Promega, G712A), GFP (to visualise GCaMP6s, 1:1000, Abcam, ab13970), NF200 (large myelinated neurons, 1:160, Sigma Aldrich, N4142), CGRP (small peptidergic neurons, 1:500, Enzo Life Sciences, CA1134) and IB4 (conjugated to Alexa Fluor 647; small non-peptidergic fibres, 1:250, Molecular Probes, I32450) overnight at room temperature. Slides were then incubated with the appropriate fluorophore-conjugated secondary antibodies (Goat anti-Chicken, Alexa Fluor 488, Invitrogen, A-11039, for GFP; Goat anti-Rabbit, Alexa Fluor 594, Invitrogen, A-11037, for CGRP and NF200, Goat anti-Mouse, Alex Fluor 647, Invitrogen, A-32728, for β-III-tubulin, all used at 1:1000 dilution) for 2 hours at room temperature. Slides were coverslipped using DAPI-containing media (Fluoromount-G with DAPI, eBioscience) and imaged with an LSM 710 laser-scanning confocal microscope (Zeiss).

### Data analysis

Drift in time-lapse recordings were corrected using NIS Elements AR 4.30.01 (Nikon, align application). Further image processing was done using Fiji/ImageJ Version 1.48v, and graphing and statistical analysis was undertaken with a combination of Microsoft Office Excel 2013, IBM SPSS Statistics 23 package and RStudio 0.99.893. All tests conducted were two-tailed and a *P* value < 0.05 was considered significant.

In order to generate traces of calcium signals from time lapse images, regions of interest (ROIs) surrounding cells were chosen using a free hand selection tool in Fiji. ROIs were chosen with minimal overlap to ensure less interference from surrounding cells.

A region of background was selected and its signal subtracted from each ROI. To generate normalised data a baseline period of fluorescence was recorded for each ROI and increases in this baseline were then calculated as ΔF/F, which was expressed as a percentage. The threshold for a positive response was taken as an average signal of 70% above baseline florescence plus 4 standard deviations (STDEV).

For tests of polymodality a high level of stringency was used. To ensure that overlapping cells were not contaminating the signal and that low levels of response were not missed, all thermally responsive cells were double checked visually for their response to pinch.

For histological analysis, to assess the success of labelling with AAV9, six to eight cells with the lowest fluorescence were selected and their average intensity plus 1 standard error of the mean (SEM) was considered the cut-off for a positive signal against which all other cells were assessed.

## Results

### Intrathecal injections of AAV9 labels a large and representative subset of neurons in L4 DRGs

In order to assess efficiency of transfection, four mice, intrathecally injected with AAV9-GCaMP6, were studied histologically 14 days post-injection. In all animals, GCaMP labelling was detected using immunohistochemistry with a GFP antibody, and all neurons identified with a β-III-tubulin antibody. A mean of 62 ± 6% of neurons were labelled with GCaMP6s following intrathecal injection (Fig. 1a). We also observed that 100% of GCaMP6s labelled cells were β-III-tubulin positive, which suggests that transduction was limited to neurons. Further, we were able to label a representative population of large myelinated (NF200 positive, 37 ± 4%), small peptidergic (CGRP positive, 34 ± 2%) and small non-peptidergic (IB4 positive, 15 ± 1%) neurons, suggesting pan-neuronal tropism (Supplementary Fig. 1).

**Figure 1.**
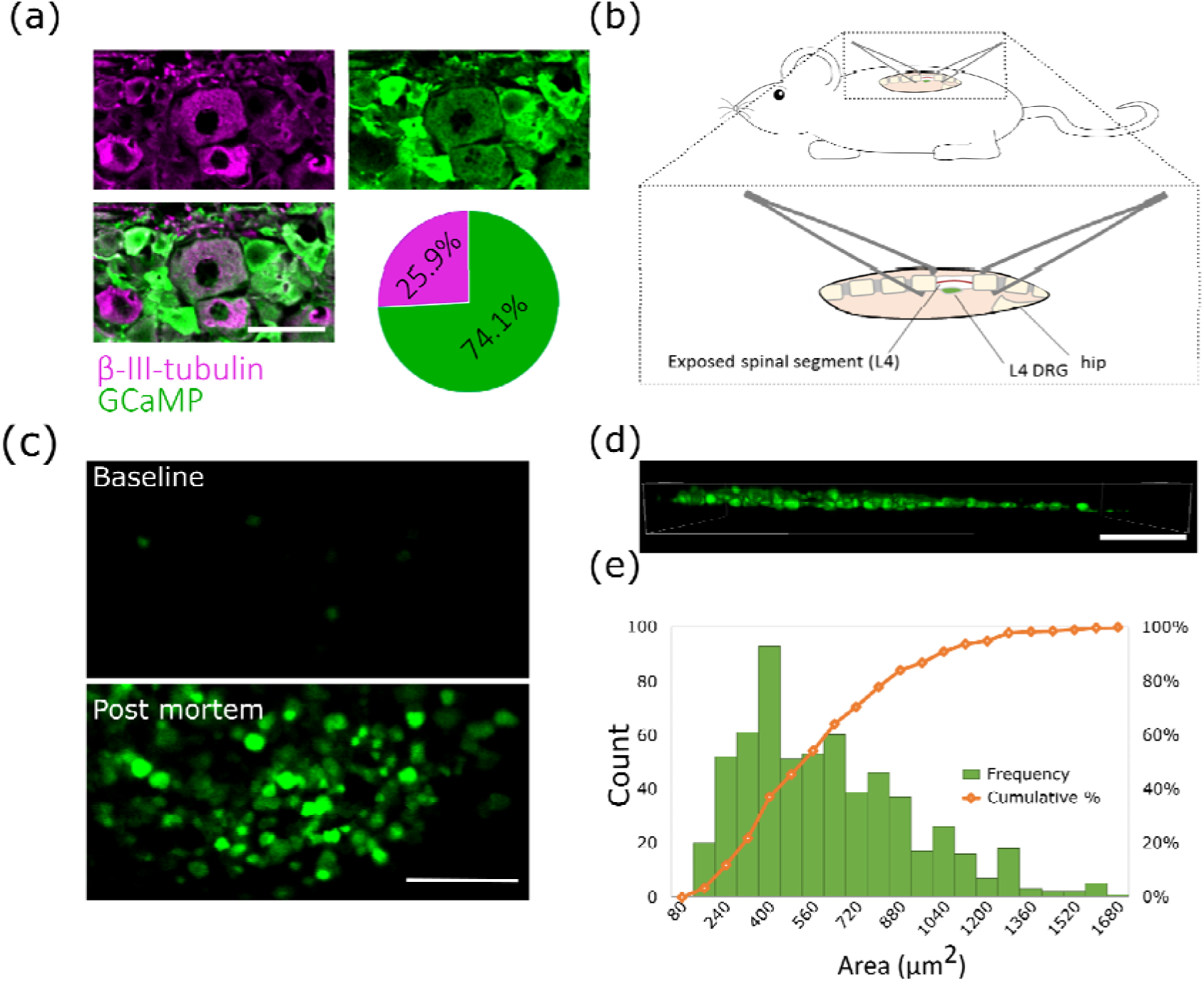
Intrathecal injections label a representative sample of DRG neurons which can be visualised using standard confocal microscopy. **(a)** Representative image and quantification of percentage of GCaMP positive cells after intrathecal injection. Scale bar = 50 μm. *n*=4 **(b)** Diagram showing the imaging set-up. The L4 DRG is exposed in a deeply anaesthetised mouse and the spinal column on either side of the exposed DRG is stabilised for *in vivo* confocal/two-photon imaging using spinal clamps attached to a custom made stage. **(c)** GCaMP6s provides a large dynamic range over which to detect signal changes *in vivo.* Very little signal is evident when the animal is unstimulated (baseline) while a large increase in signal strength is evident following intracellular calcium accumulation >30 minutes after death *(post mortem).* Scale bar = 200 μm. **(d)** A 3D reconstruction of a z stack reveals that an approximately one-cell-thick layer is being assessed using this *in vivo* imaging technique. Scale bar = 200 μm. **(e)** Frequency histogram and cumulative sum percentage of differentially sized cells shows a skewed distribution with a larger percentage of smaller cells. *n*=3456 cells in *n*=13 mice.

### *In vivo* imaging of GCaMP6-labelled cells: confocal vs. two-photon microscopy

We exposed and visualised the L4 DRG *in vivo* and were able to detect consistent labelling as early as 2 weeks after AAV9-GCaMP6 injection. The primary stability for imaging was provided by clamping the spinous process above and below the L4 vertebra with forceps (Fig. 1b). Rotation of the mouse and spinal cord position in the horizontal axis optimised the orientation of the DRG towards the optical plane of the objective and ensured maximal focus of DRGs during *in vivo* imaging.

We found that confocal microscopy with a maximally opened pinhole offered several advantages over multiphoton microscopy in our preparation. Confocal imaging provided more available signal, including from out-of-focus cells (Supplementary Fig. 2a). As a result, more information could be extracted from a larger number of cells (Supplementary Fig. 2a & b). In addition, the greater optical slice thickness reduced the negative effects of motion and provided more stable traces of fluorescence over time (Supplementary Fig. 2b), which is particularly important when stimulation induces some movement (e.g. with electrical stimulation, see below). Furthermore, the increased signal (compared with the thinner z slices available with two-photon imaging) required lower laser strengths and offered faster acquisition. Using this technique, we were able to image at more than 16 Hz for over an hour without visible photo-damage. These relative benefits of confocal microscopy in this preparation may be enhanced by the absence of a coverslip and the curved shape of the DRG which result in a less regular air/sample interface and more out of focus tissue.

Using the confocal approach outlined above it was possible to assess our minimal and maximal imaging range through comparison of cellular fluorescence at baseline (without any stimulation) and following cellular calcium accumulation more than 30 minutes after death (Fig. 1c). This revealed a large dynamic range over which we could visualise changes in intracellular calcium. A 3-dimensional (3D) reconstruction of a z-stack of the imaged DRG when viewed from the side revealed an approximately one-cell-thick layer over which images were acquired using this confocal technique (Fig. 1d).

Using a 10X objective lens, we were typically able to image over 200 neurons simultaneously per DRG. Estimation of cell size shows an expected unimodal distribution with a majority of small diameter cells and a tail of fewer, large diameter somas (Fig. 1e).

In the absence of stimulation most cells had stable fluorescence with a STDEV of around 7%. Only 79 of 10620 cells (~5%) showed significant spontaneous fluctuations, which are likely to represent ongoing activity (from comparisons with evoked responses, see below).

### Assessing the sensitivity of GCaMP6s sensory neuron imaging with electrical stimulation of the sciatic nerve

Due to the novelty of our technique it was necessary to establish the relationship between action potential firing and calcium transients in the DRG. We took advantage of the fact that action potentials can be elicited in the DRG with a precisely controlled frequency and temporal pattern via electrical stimulation of peripheral nerves.

We used direct electrical stimulation of the sciatic nerve (which contains axons of the large majority of L4 DRG cells) to induce activity in either large myelinated sensory neurons (A-fibres) or large and small afferents (A and C-fibres) at different frequencies (Supplementary Video 1). Each stimulus induces an action potential that propagates into the DRG and depolarises the cell body.

We found that, with A-fibre strength stimulation, a subset of electrically responsive L4 DRG neurons (48%) showed increased fluorescence (Fig. 2a-c). The average magnitude of the A-fibre maximal response at 20 Hz was 654 ± 974% of basal fluorescence. Supramaximal sciatic nerve stimulation activated the same cells, but additionally recruited the large majority of remaining labelled cells in the L4 DRG. These additional cells, only responding to high intensity stimulation, were presumed to be cells with C-fibre axons. They showed an average magnitude of 868 ± 765% of basal fluorescence at their maximal response during 20 Hz stimulation. In both A-fibres and C-fibres the fluorescence intensity increased with stimulus frequency (Fig. 2b & c), suggesting that GCaMP6 fluorescence intensity provides a useful proxy for frequency of action potential firing.

**Figure 2.**
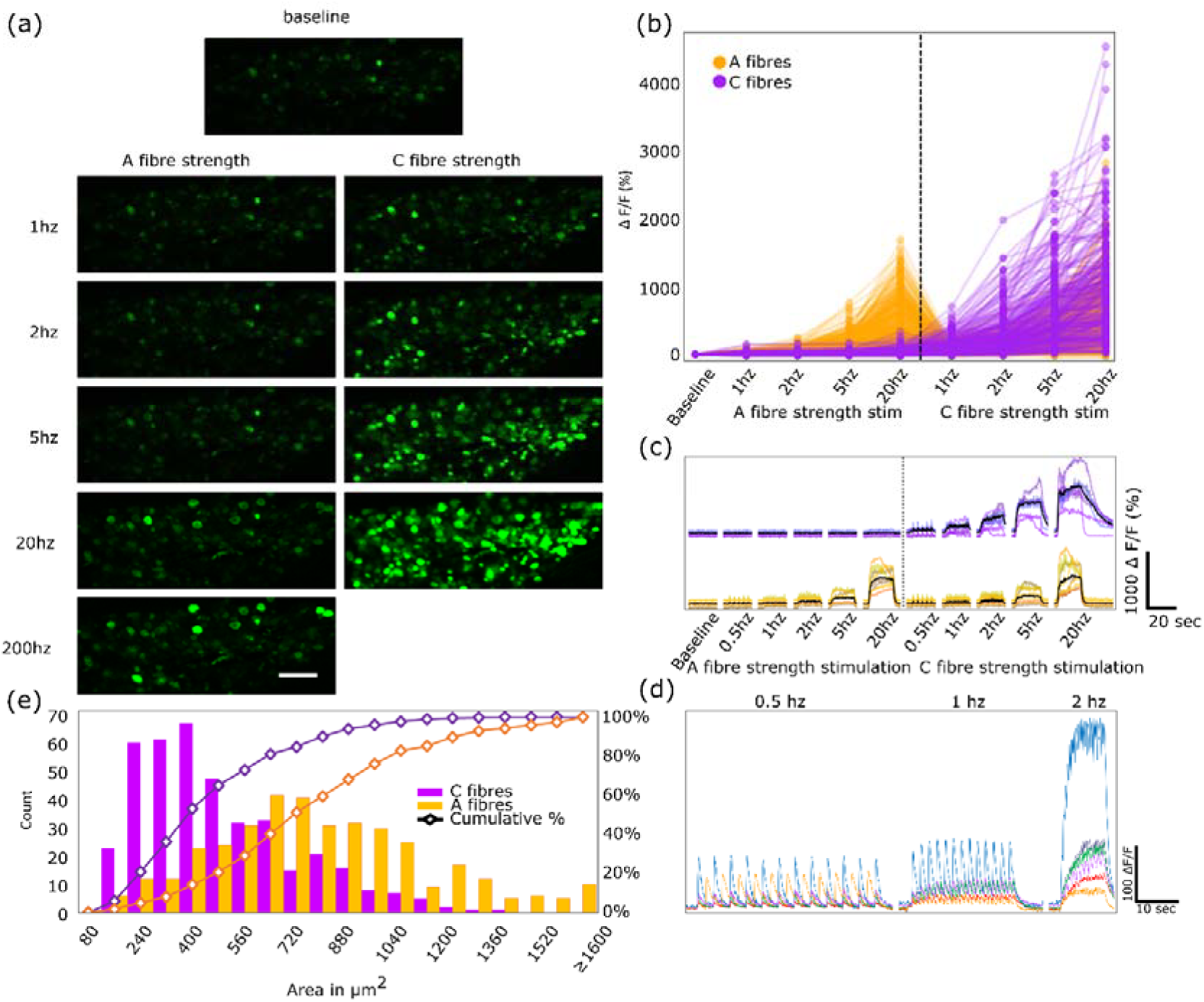
Electrical stimulation of the sciatic nerve leads to an increase in calcium signals in the DRG in an intensity and frequency dependent manner. **(a)** Images of DRG cell bodies during direct stimulation of the sciatic nerve. Recruitment of a greater number of cells occurs at higher intensities and frequencies. 200 Hz stimulation was omitted from C-fibre strength stimulation to avoid damage to the nerve. Scale bar = 200 μm. **(b)** Traces of individual cells responding to electrical stimulation. Yellow lines represent cells that respond to both A- and C-fibre strength stimulation and purple lines represent cells that respond to C-fibre strength stimuli only. Data points displayed at averaged intensity over the period of stimulation. *n*=774 cells in *n*=6 mice. Two outliers were removed from this graph for scaling purposes but were retained for analysis **(c)** Representative traces of GCaMP fluorescence during electrical stimulation. Electrical stimulation at A- and C-fibre strength resulted in activation of distinct subsets of cells in a frequency dependent manner. Black lines represent averaged data. **(d)** Zoomed representative traces of GCaMP fluorescence during stimulation at C-fibre strength at low frequencies (0.5h Hz, 1 Hz and 2 Hz) and high frame acquisition rate (8 Hz). Single action potentials generated by single electrical pulses to the sciatic nerve could be detected in the DRG. Each trace represents fluorescence from a single cell. **(e)** Size distribution of cells that respond to A-fibre strength stimulation (yellow) compared to cells that respond to C-fibre strength stimuli (purple) and the cumulative sum of their sizes. Cells responsive to A-fibre strength stimulation are significantly larger compared to cells which only respond to C-fibre strength stimuli: two-sample Kolmogorov-Smirnov test, *n*=776 cells in *n*=6 mice, *P*<0.001.

When the stimulation frequency was sufficiently low and the image acquisition rate high, a fluorescence signal was seen in response to each arriving impulse in the majority of neurons, revealing sufficient sensitivity to detect single action potentials (Fig. 2c & d). To ensure that only action potentials generated through stimulation of the sciatic nerve were recorded, the stimulated nerve was transected distally to the stimulation site (Fig. 2d). This procedure ensured an absence of contaminating spontaneous peripheral inputs, confirmed by the lack of response to paw pinching. The magnitude of this unitary response varied greatly, in some cases exceeding 100% of baseline fluorescence, typically had a rise time of around 200 milliseconds, and an exponential decay with a time constant of approximately 700 milliseconds.

As expected, the size distribution of cells responding to A-fibre stimulation was on average larger than those cells recruited only by C-fibre strength stimulation (Fig. 2e).

### Assessing polymodality in primary afferents

In order to assess the level of polymodality, all thermally responsive cells were assessed for their responses to the opposing thermal modality (i.e. cold responding cells that also respond to heat and vice versa) as well as their mechanical sensitivity.

A small proportion of heat-responding cells (~4%) also responded to decreases in temperature, while a larger proportion of cold responding cells (~40%) also encoded an increase in temperature (Fig 3a & b). However, due to the very small number of cold responsive neurons these percentages varied greatly between experiments. Around 55% of temperature responsive cells were also responsive to pinch, in line with previous evidence revealing a subset of multimodal temperature sensitive C-fibres (Fig. 3a - c).

**Figure 3.**
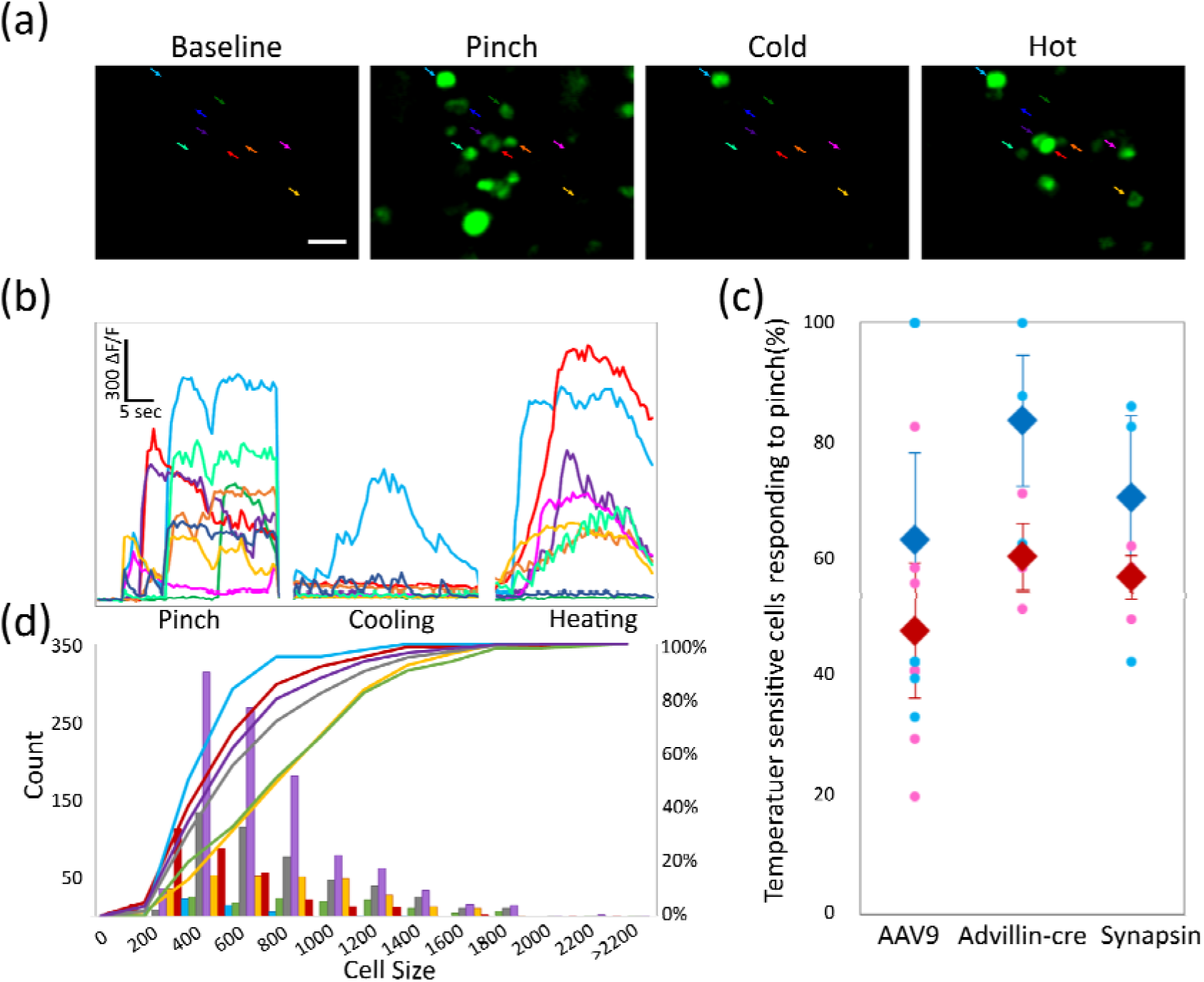
Polymodality is common in primary afferent nociceptors. **(a)** Example images of DRG cells responding to mechanical and thermal stimulation of the ipsilateral plantar paw. Scale bar = 50μm. **(b)** Colour coded example traces of cells highlighted in **(a). (c)** Percentage of thermally sensitive DRG cells responding also to noxious mechanical stimulation. Similar percentages of polymodal (thermally and mechanically sensitive cells) were detected when using AAV9 to deliver GCaMP through intrathecal injections (*n*=6) or when expressing GCaMP transgenically only in DRG neurons through GCaMP floxed mice expressing cre under the Advillin promoter (peripheral neurons) (*n*=3) or when expressing GCaMP in all neurons through the Snap25 promoter (*n*=3). Total *n*=1138 cells. Blue data points show the percentage of cold sensitive neurons responding to pinch while red data points indicate the percentage of hot responding cells also responding to pinch. Diamonds display the mean ± SEM, circles indicate single mice. **(d)** Cell size distribution of cells that respond to increases/decreases in temperatures vs cells that respond to A-fibre strength stimulation, C-fibre strength stimulation, brush and pinch and the cumulative sum of their sizes. Cells responsive to A-fibre strength stimulation, C-fibre strength stimulation, brush and pinch are significantly larger compared to cells which respond to changes in temperature: two-sample Kolmogorov-Smirnov test, total *n*=1166 cells in *n*=4 mice, *p*<0.001 for all comparisons against temperature sensitive cells, except temperature responsive cells vs C-fibre stimulation where *p*<0.02.

A size distribution showed that thermosensitive DRG cells were significantly smaller than cells of the same cohort of animals that responded to stimulation by brush and A-fibre strength stimulation (Fig. 3d). Indeed cold-sensitive cells represented the smallest pool of cells with pinch-sensitive cells and those responding to C-fibre strength stimulation being of average size, in agreement with the non-discriminative nature of these stimuli.

To ensure that our labelling technique did not lead to a selection bias, the responsiveness of temperature sensitive cells to mechanical stimulation (pinch) was also assessed in two different lines of GCaMP6 transgenic mice (Fig 3c). Despite the use of three different approaches to label DRG neurons (intrathecal injections of AAV9 carrying the GCaMP6 transgene, vs two types of transgenic animals expressing GCaMP6 constitutively in all neurons or conditionally only in peripheral neurons), the polymodality of nociceptors was maintained (Fig 3c).

### Assessing the effect of an intraplantar injection of formalin on prolonged DRG activity

The formalin test is very widely used in pain research but the mechanisms underpinning the responses are not completely understood. To address this question we injected 20μl formalin (1.85% in saline) or vehicle into the plantar surface of the hind paw, while simultaneously imaging the ipsilateral DRG.

An injection of saline produced, as expected, a brief burst of activity, presumably from the pressure associated with the volume increase in the paw. In contrast, the formalin injection induced a significantly greater response, much of it with a delayed activation (Fig. 4a & b) with some cells showing brief and sometimes repeated bursts of activity a few minutes after a formalin injection (Fig. 4a, c & d). Indeed, the composite activity of DRG cells mirrored pain behaviour up to the second peak (Fig. 4a & b). However, following the interphase, the average response of DRG cells no longer corresponded to the behavioural pattern and, instead, minimal activity, similar to saline, was observed.

**Figure 4.**
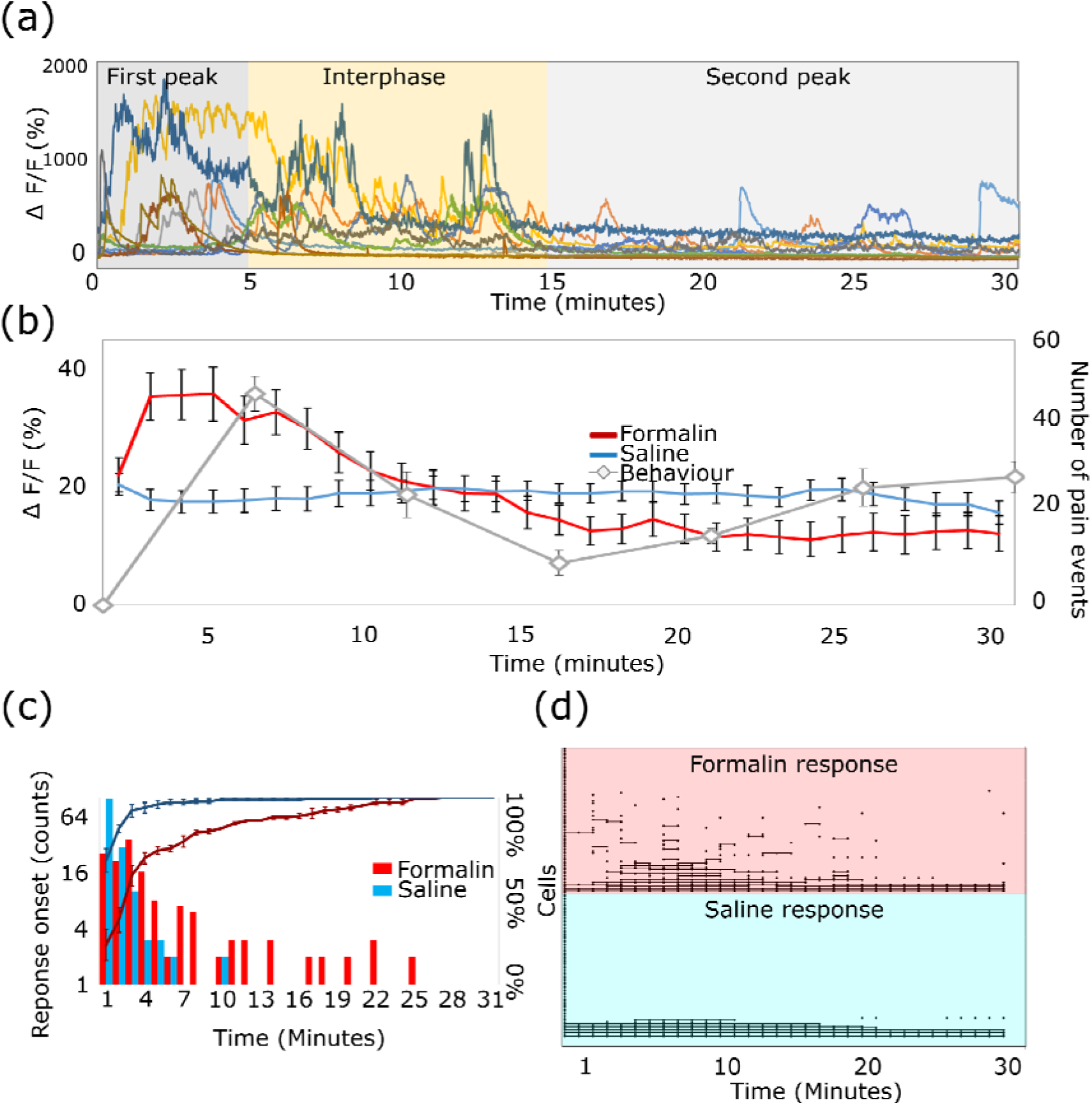
DRGs play a defining role in the generation of pain behaviour in the formalin test. **(a)** Example traces of GCaMP fluorescence 0-30 minutes after intraplantar injection of formalin (1.85%). **(b)** Average response of DRG cells (*n*=666 cells in *n*=5 mice) and behavioural response in mice (n=8) after injection of formalin and saline into the plantar surface of the hind paw. Cellular response was averaged over one minute and across cells. Pain behaviour was assessed as the number of pain events displayed by the mouse in 5 minute increments. All data displayed as means ± SEM. Split plot ANOVA (no equality of variance assumed), interaction of treatment with time: *P*<0.001. **(c)** Onset of first response after saline/formalin injection. Bars represent the number of cells that showed their first response at different times following injection. The lines represent the percentage of cells which have already responded. Two-sample Kolmogorov-Smirnov test, *n*=666 cells in *n*=5 mice, *P*<0.001. **(h)** Sample binary responses of cells to an injection of saline/formalin. Cells are stacked on the Y-axis and each point at time 0 indicates a separate cell. A response is defined as a signal more than 70% above baseline + 4 STDEV and is plotted as a black point. Connecting lines are drawn if continuous firing occurred over two or more time points. The same cells were plotted in response to saline and formalin.

### UVB irradiation leads to an increased sensitivity of thermally responsive neurons

UVB irradiation is used as a model of inflammatory pain in humans and animals. We assessed the consequences of this treatment on the responsiveness of sensory neurons. We found that UVB inflammation consistently led to a striking increase in the responsiveness of temperature sensitive neurons (Fig. 5a & b). Indeed, warming of the injured paw lead to striking increases in the responses of affected primary afferents which now revealed a linear increase in response with increases in temperature. This is compared with much lower levels of activation after heating of the uninjured paw. Here low levels of heating have very limited effects on neuronal activity and much higher levels of stimulation are needed to achieve significant increases in neuronal responses (Fig. 5a & b). The sensitisation of the peripheral nervous system was not limited to heating but also included enhanced sensitivity towards cooling of the injured paw. Even small decreases in temperature (e.g. 27°C) resulted in much greater activation of responding cells during sterile inflammation as compared with responses in healthy control animals (Fig. 5b). In addition, responses to both noxious (pinch) and innocuous (brush) mechanical stimuli were significantly enhanced after UVB irradiation (Fig. 5c, see Supplementary Fig 3. and Supplementary Video 2 for sample traces of mechanical stimulation).

**Figure 5.**
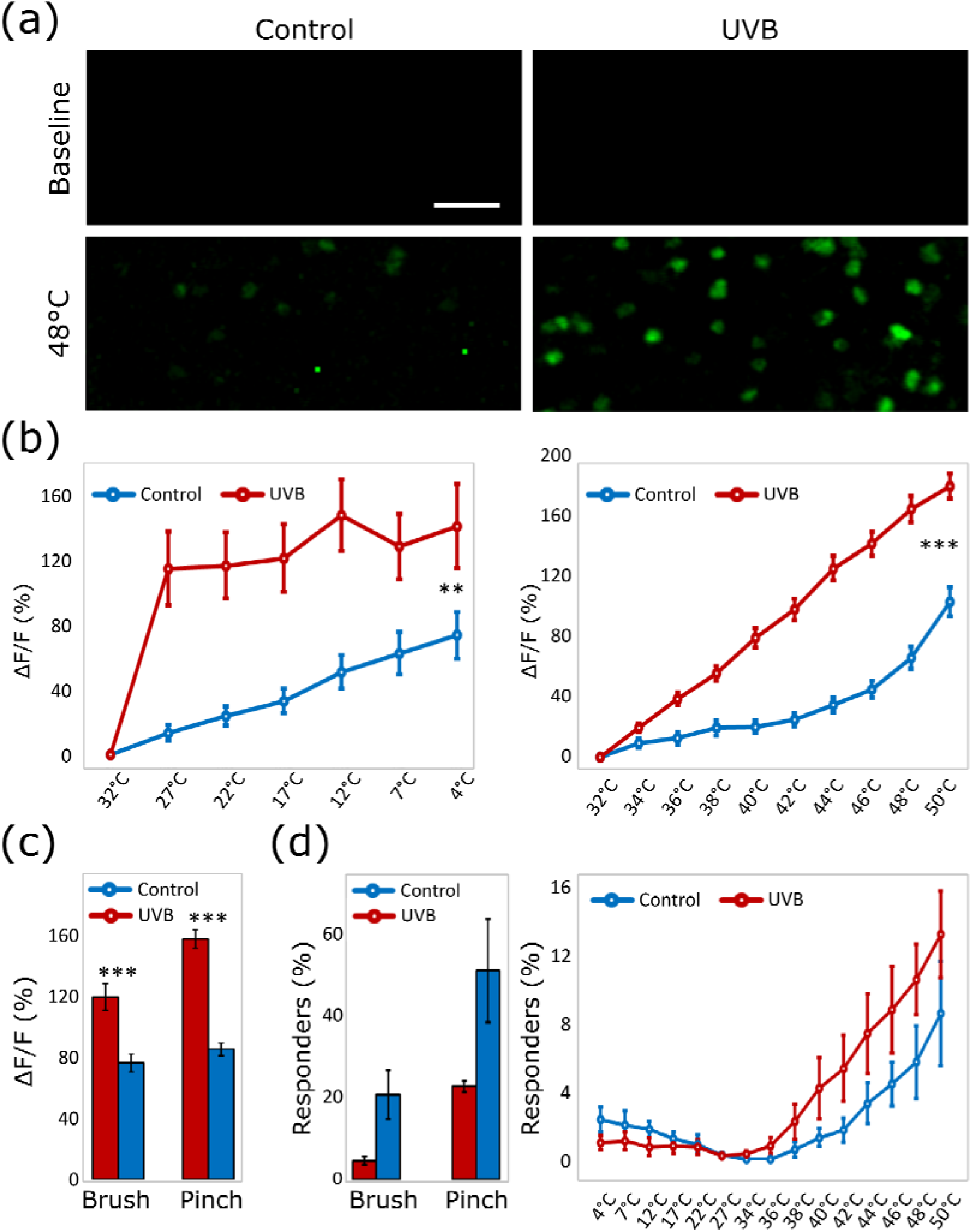
UVB irradiation increases the responsiveness of peripheral neurons. **(a)** Representative images of DRG neurons in mice stimulated with 32°C (baseline) and 48°C on the ipsilateral plantar surface, 48hrs after UVB irradiation or control anaesthesia. Scale bar = 100μm. **(b)** The response intensity of neurons stimulated thermally 48hrs after UVB irradiation or control anaesthesia. The response of neurons to both warming (n=393 cells) and cooling (n=71 cells) of the ipsilateral paw is significantly greater in UVB irradiated animals compared to controls (between groups difference in a split-plot ANOVA, ** = p<0.005, *** = p<0.001). **(c)** Response intensities of neurons stimulated mechanically (brush: n=222 cells; pinch: n=787 cells) 48 hrs after UVB irradiation are significantly greater as compared to neurons in sham irradiated mice (independent sample t-test, equal variances not assumed, *** = p<0.001). **(d)** Percentage of cells responding to different types of peripheral stimulation does not vary significantly after UVB irradiation vs sham anaesthesia. For all experiments data displayed as means ± SEM, n=7 mice (3 control, 4 UVB).

Interestingly UVB irradiation did not lead to an increase in the percentage of cells responding to thermal or mechanical stimulation of the irradiated paw, when compared to control anaesthetised mice (Fig. 5d).

## Discussion

Here, we use a new technique for the visualisation and study of hundreds of peripheral cells simultaneously *in vivo.* Using advanced genetically encoded calcium indicators, together with standard confocal microscopy, we were able to assess population responses of DRG neurons to electrical, thermal, mechanical and chemical stimuli, providing new insights into the processing of sensation and nociception in the peripheral nervous system during health and injury. While there have been a few reports using similar approaches, this is the first to report on large populations (typically more than 200 cells) in each preparation acquired at a sub second framerate.

The exposure and stabilisation of the DRG involved a laterally extended laminectomy. Due to the encasing bony structure of the vertebrae, sufficient stabilisation only required two vertebral clamps on either side of the exposure. However, the orientation of the DRG around the spinal cord meant that aligning the entire structure into one focal plane was challenging. As a result, confocal microscopy with an open pinhole was considered beneficial over two-photon microscopy in this instance. This approach is complementary to other methods, using both multi- and single-photon microscopes, aimed at increasing the acquisition of information in space while maintaining temporal resolution. Such methods include for example the use of a piezoelectric device (Callamaras and Parker, 1999), Bessel beams (Thériault et al., 2014), acousto-optical deflectors (Reddy and Saggau, 2005), multi-spot excitation (Bewersdorf et al., 1998; Kurtz et al., 2006), as well as various adaptations of scanning patterns (Göbel et al., 2007; Lillis et al., 2008). While our approach loses some of the added benefits often associated with two-photon microscopy, including greater penetration depth and reduced photo damage, it requires no modification of the microscope or software and provides a large dynamic range over which to detect changes in calcium signal, with a favourable signal-to-noise ratio and reduced effects of movement artefacts. In addition, the signal remained stable across long imaging sessions (>2hrs) with no detected bleaching.

The functionality of our system was validated by electrical stimulation of the sciatic nerve, activating DRG neurons in a predictable and reproducible manner (Fig. 2). The results of our experiments indicate that GCaMP6 fluorescence in DRG neurons is a sensitive reflection of neuronal activity, with good temporal resolution. Interestingly our approach resulted in much greater numbers of neurons detected at much faster rates, compared to previously published works (Emery et al., 2016; Kim et al., 2016; Smith-Edwards et al., 2016). As expected, the majority of L4 DRG neurons responded to stimulation in a frequency dependent manner: low threshold electrical stimuli recruited fewer cells in the DRG, and these tended to have larger DRG somata. In contrast, high threshold stimulation activated both the same large cells and additionally many more (typically small diameter) neurons. Moreover, the GCaMP6 reporter was sensitive enough to detect calcium transients in the DRG elicited by single action potentials generated in the periphery.

At stimulation frequencies above 1 Hz, temporal summation of the calcium transient meant that the fluorescence intensity also served as a proxy for the rate of action potential firing. The magnitude of calcium transients for a given level of activity varied between cells. Some of this presumably represents variation in expression levels of GCaMP6s in different cells, but it may also represent differences in calcium handling by different cell types (Rausell et al., 1992; Malmberg and Yaksh, 1994; Westenbroek et al., 1998; Pirec et al., 2001; Simons et al., 2009).

Having established the sensitivity and specificity of the technique, we next investigated several aspects of the neurobiology underpinning sensory perception. Due to the novelty of the method and several technical challenges it poses, it is essential to carefully examine novel claims that contrast traditionally held views. Indeed, one such claim suggests that polymodality, traditionally seen using microneurography, electrophysiology, multiple lines of *in vitro* evidence, and now also *in vivo* GCaMP imaging, was due to inflammation introduced through the use of invasive techniques (Emery et al., 2016). However, our data shows that this is unlikely to be an adequate explanation as the use of three different labelling techniques (including transgenic expression of GCaMP similar to that used by Emery et al., 2016) resulted in similar levels of polymodality. It is possible that in the evolving stages of peripheral nervous system imaging the small numbers of recorded cells reported in the previous studies can potentially skew results in unexpected ways as applications can still miss one of the most essential benefits of peripheral imaging which is the collection of large sets of unbiased observations.

With large number of cells visualised with a good temporal resolution however, this technique can provide a perfect system with which to study the involvement of the peripheral nervous system in several acute and chronic pain conditions. The formalin test is one of the most widely used models for the study of pain, particularly in the pharmaceutical industry. Controversy still exists as to its mechanism of action, particularly relating to the relative role of the peripheral vs central nervous system. It has been variably suggested that the second phase is generated directly by afferent activity (McCall et al., 1996) or by sensitisation of the spinal dorsal horns (Coderre et al., 1990, 1993; Malmberg and Yaksh, 1992), or even supraspinal structures (Vaccarino and Melzack, 1989). The ability to visualise hundreds of DRG cells simultaneously was perfectly suited to investigate the relative role of the peripheral network in the generation of the formalin response.

Our results show high levels of DRG activation soon after an injection of formalin with a peak activity 5 minutes post-injection. The cellular response was enhanced and delayed relative to saline, in line with previous work (Puig and Sorkin, 1996). We found a range of responses after formalin injection. A small number of cells, typically firing for long periods of time post-injection, were likely responding to distention associated with the injection, since similar responses were seen after saline injections. Beyond these, there were many cells that were activated selectively by formalin. Some were activated soon after the injection, while others showed a more delayed response – either in the form of a single ‘burst’ or of multiple cycles of activity across the observed period. Crucially, in contrast to previous electrophysiological data (McCall et al., 1996; Henry et al., 1999), our recorded cells did not seem, in aggregate, to constitute a ‘second phase’ of inputs to the spinal cord. Rather, our population studies show that overall DRG activity, during the first peak and interphase, corresponds well with the observed pain behaviour, but then diverges around the second peak. In light of this, our data suggest that afferent activity is capable of driving the first phase in behaviour while simultaneously sensitising the CNS. Due to this sensitisation, low levels of peripheral activity are sufficient to induce a second phase, which is sensitive to blockers of central sensitisation during the first phase, but not ablated by local peripheral anaesthesia during the second phase (Dickenson and Sullivan, 1987; Coderre et al., 1990; Murray et al., 1991; Yamamoto and Yaksh, 1992).

The divergence between our findings and classical electrophyisological investigations may have several reasons: generally, electrophysiology reports that neuronal activity during the second phase is considerably reduced and may therefore not be sufficient to induce significant calcium fluctuations in DRG neurons. Furthermore, *In vivo* imaging can easily capture the large variation in responses of different cell types and their dependence on the distance of the injection from the receptive field of the responding cell (Heapy et al., 1987; Puig and Sorkin, 1996). Conversely, electrophysiology is limited to small cell numbers, and therefore cannot easily cover a representative selection of cells. This benefit applies to numerous other questions that are traditionally studied using electrophysiological measures, as *in vivo* imaging can avoid selection biases, for example, against cells that are mechanically insensitive or even silent (Shoham et al., 2006). Such an unbiased approach can equally be applied to other questions and sensory modalities, including the study of itch or sensitisation/priming during physiological and also pathological states.

Indeed, one interesting and highly translatable model of persistent pain is UVB induced sterile inflammation. UVB irradiation leads to consistent swelling and hyperalgesia 48 hrs after injury. Here we were able to directly assess the involvement of the peripheral nervous system in the resulting hyperalgesia (Fig. 5). Indeed, we were able to visualise significant increases in sensitivity to peripheral heating and peripheral mechanical stimulation (both noxious and innocuous) as reported previously using electrophysiological methods (Bishop et al., 2010). Additionally, we also demonstrate for the first time, to the best of our knowledge, the involvement of the peripheral nervous system in UVB induced cold sensitivity. Indeed, we show that even small decreases in temperature had much more profound effects on peripheral responses after UVB irradiation as compared with responses from control animals. Interestingly no significant differences were observed in the percentage of neurons responding to mechanical of thermal stimuli. This suggests limited unmasking of silent cells with this model of sterile inflammation, and instead implicates mainly an increased activation of responding cells.

It should be noted that *in vivo* imaging provides a significant advantage over traditional electrophysiological techniques in questions of cold processing. The limited number of cold sensitive primary afferents provides a practical barrier for the assessment of changes in peripheral cold processing using stochastic single cell approaches. Instead, the ability to sample hundreds of cells simultaneously, as shown here, provides an excellent opportunity to investigate populations of rare neurons, including, but not limited to, cold sensitive afferents.

In conclusion, we significantly enhanced the spatial and temporal sensitivity of a novel technique applied to the large scale study of the peripheral nervous system. Using this technique we were able to confirm the traditionally held, but recently challenged, view of polymodality in the peripheral nervous system. Additionally we revealed new roles for peripheral processing in acute and persistent pain states.

## Acknowledgement

The Genetically-Encoded Neuronal Indicator and Effector (GENIE) Project and the Janelia Research Campus of the Howard Hughes Medical Institute has generously allowed this material to be distributed with the understanding that requesting investigators need to acknowledge the GENIE Program and the Janelia Research Campus in any publication in which the material was used, specifically Vivek Jayaraman, Ph.D., Douglas S. Kim, Ph.D., Loren L. Looger, Ph.D., Karel Svoboda, Ph.D. from the GENIE Project, Janelia Research Campus, Howard Hughes Medical Institute.

Thank you to Professor Peter McNaughton and Tamara Buijs for the use of Snap25-2A-GCaMP6s-D mice. Also thank you to Tatum Cummins and Dr Franziska Denk for valuable comments on the MS and Dr Tommaso Tufarelli and Dr Sarah Chisholm for expert statistical advice.

---

**Supplementary Figure 1.**
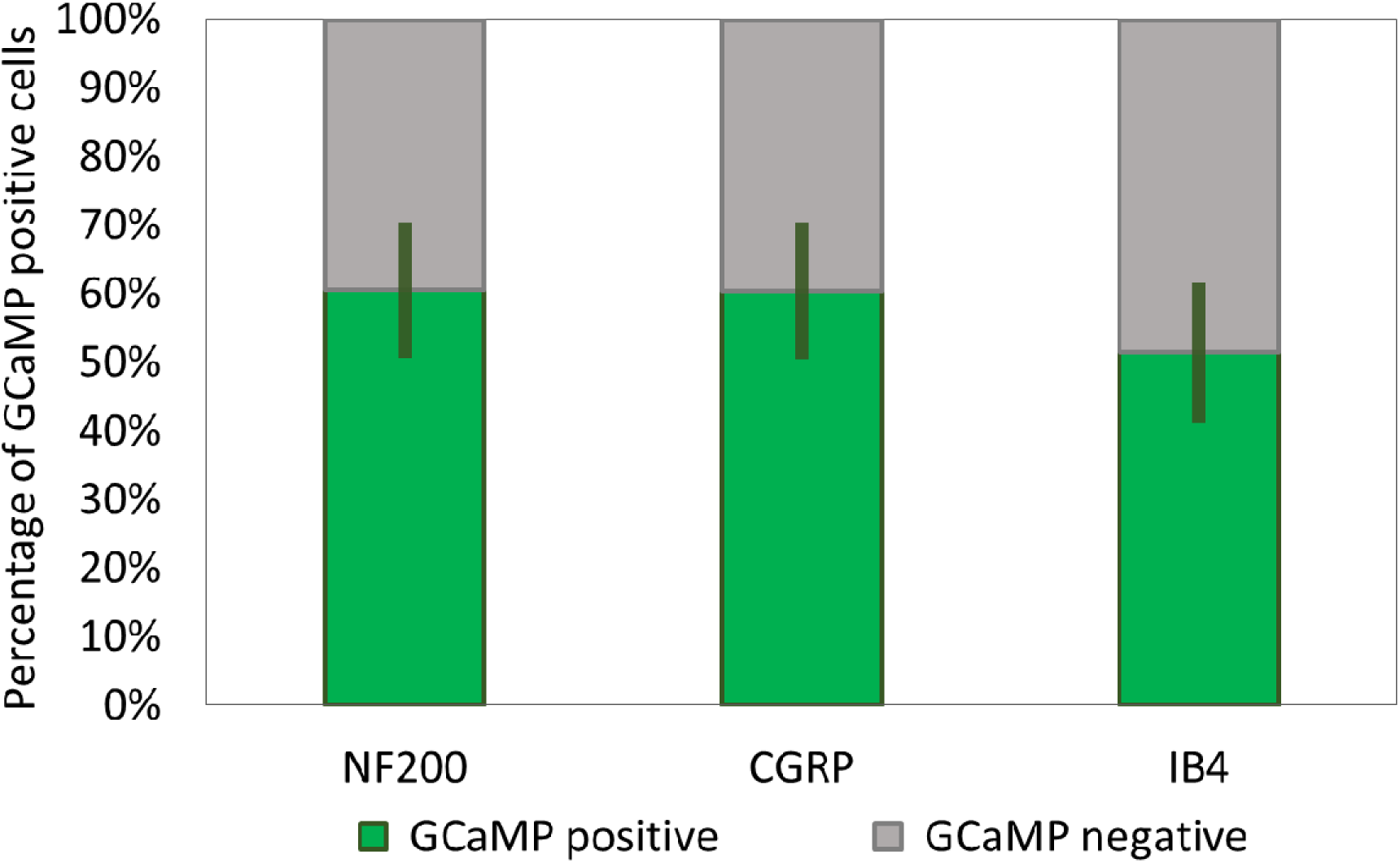
Proportion of GCaMP labelled cells. Quantification of GCaMP positive cells as a proportion of all NF200, CGRP and IB4 immunoreactive cells. Data displayed as mean ± SEM. *n*= 4 mice.

**Supplementary Figure 2.**
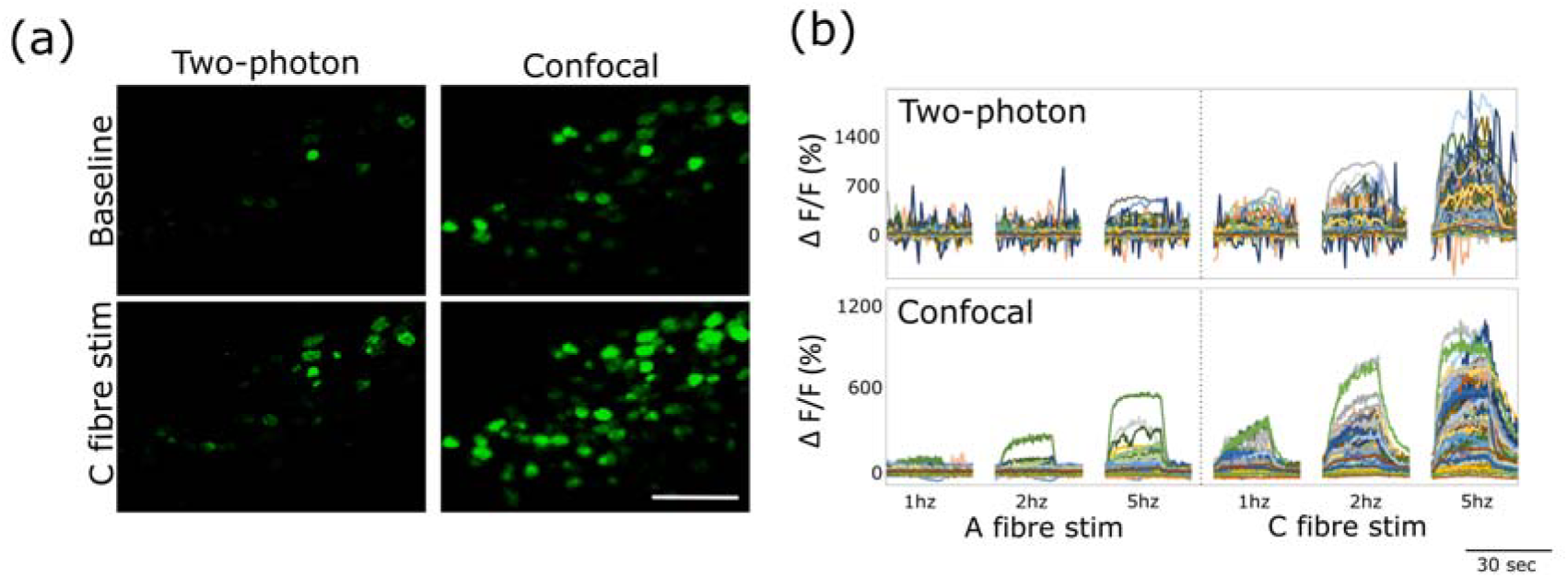
The use of confocal microscopy vs two photon microscopy. **(a)** Confocal microscopy with an open pinhole reveals cellular GCaMP signal which is lost in two-photon acquisition mode. Due to the more restricted slice thickness of multiphoton microscopy (left panel) fewer cells are visible in the same field of view compared to when confocal microscopy is used with an open pinhole (right panel). This is particularly obvious during stimulation of the sciatic nerve (lower panel). Scale bar = 200 μm. **(b)** Fluorescence traces of **(a).** The sciatic nerve was stimulated at 1, 2 and 5 Hz at both A- and C-fibre strengths to achieve a representative view of different response amplitudes both during confocal and multiphoton acquisition. In this preparation a cleaner signal is generated when confocal microscopy with an open pinhole is used compared to two-photon microscopy.

**Supplementary Figure 3.**
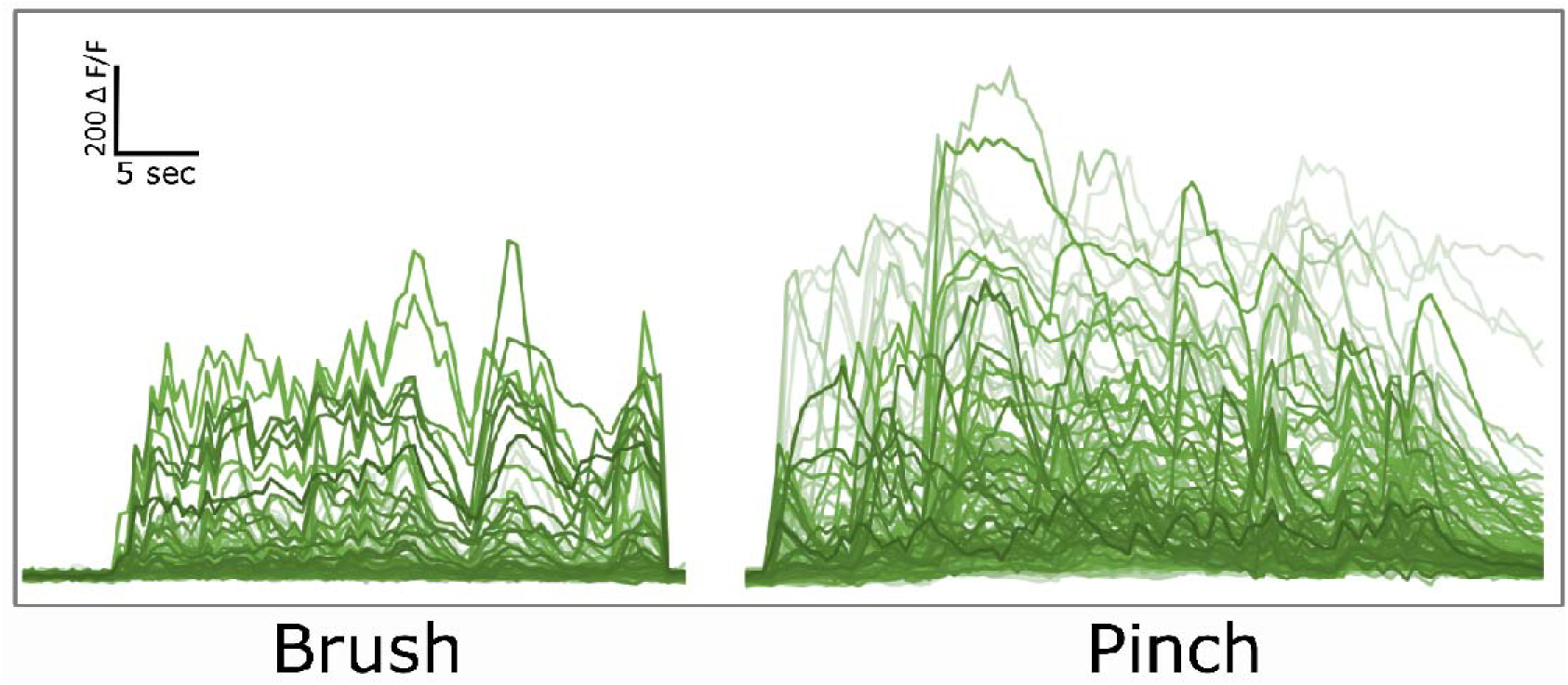
Example traces of cells responding to repeated mechanical stimulation (brush and pinch). Traces taken from Supplementary video 2.

## References

Barretto RPJ, Gillis-Smith S, Chandrashekar J, Yarmolinsky DA, Schnitzer MJ, Ryba NJP, Zuker CS (2014) The neural representation of taste quality at the periphery. Nature 517:373–376.

Bewersdorf J, Pick R, Hell SW (1998) Multifocal multiphoton microscopy. Opt Lett 23:655.

Bishop T, Marchand F, Young AR, Lewin GR, McMahon SB (2010) Ultraviolet-B-induced mechanical hyperalgesia: A role for peripheral sensitisation. Pain 150:141–152.

Callamaras N, Parker I (1999) Construction of a confocal microscope for real-time x-y and x-z imaging. Cell Calcium 26:271–279.

Chen Q, Cichon J, Wang W, Qiu L, Lee S-JR, Campbell NR, Destefino N, Goard MJ, Fu Z, Yasuda R, Looger LL, Arenkiel BR, Gan W-B, Feng G (2012) Imaging neural activity using Thy1-GCaMP transgenic mice. Neuron 76:297–308.

Chen T-W, Wardill TJ, Sun Y, Pulver SR, Renninger SL, Baohan A, Schreiter ER, Kerr RA, Orger MB, Jayaraman V, Looger LL, Svoboda K, Kim DS (2013) Ultrasensitive fluorescent proteins for imaging neuronal activity. Nature 499:295–300.

Coderre TJ, Katz J, Vaccarino AL, Melzack R (1993) Contribution of central neuroplasticity to pathological pain: review of clinical and experimental evidence. Pain 52:259–285.

Coderre TJ, Vaccarino AL, Melzack R (1990) Central nervous system plasticity in the tonic pain response to subcutaneous formalin injection.

Dana H, Chen T-W, Hu A, Shields BC, Guo C, Looger LL, Kim DS, Svoboda K (2014) Thy1-GCaMP6 transgenic mice for neuronal population imaging in vivo. PLoS One 9:el08697.

Dickenson AH, Sullivan AF (1987) Subcutaneous formalin-induced activity of dorsal horn neurones in the rat: differential response to an intrathecal opiate administered pre or post formalin. Pain 30:349–360.

Dombeck DA, Khabbaz AN, Collman F, Adelman TL, Tank DW (2007) Imaging large-scale neural activity with cellular resolution in awake, mobile mice. Neuron 56:43–57.

Emery EC, Luiz AP, Sikandar S, Magnúsdóttir R, Dong X, Wood JN (2016) In vivo characterization of distinct modality-specific subsets of somatosensory neurons using GCaMP. Sci Adv 2.

Flusberg BA, Nimmerjahn A, Cocker ED, Mukamel EA, Barretto RPJ, Ko TH, Burns LD, Jung JC, Schnitzer MJ (2008) High-speed, miniaturized fluorescence microscopy in freely moving mice. Nat Methods 5:935–938.

Ghosh KK, Burns LD, Cocker ED, Nimmerjahn A, Ziv Y, Gamal A El, Schnitzer MJ (2011) Miniaturized integration of a fluorescence microscope. Nat Methods 8:871–878.

Göbel W, Kampa BM, Helmchen F (2007) Imaging cellular network dynamics in three dimensions using fast 3D laser scanning. Nat Methods 4:73–79.

Heapy C, Jamieson A, Russell N (1987) Afferent C-fiber and A-delta activity in models of inflammation. Br J Pharmacol 90:23.

Henry JL, Yashpal K, Pitcher GM, Coderre TJ (1999) Physiological evidence that the ‘interphase’ in the formalin test is due to active inhibition. Pain 82:57–63.

Johannssen HC, Helmchen F (2010) In vivo Ca2+ imaging of dorsal horn neuronal populations in mouse spinal cord. J Physiol 588:3397–3402.

Kandel ER, Schwartz JH, Jessell TM, Siegelbaum SA, Hudspeth AJ (2013) Principles of neural science, 5th ed. McGraw-Hill.

Kim YS, Anderson M, Park K, Guan Y, Spray DC, Dong X, Shin Kim Y, Zheng Q, Agarwal A, Gong C, Young L, He S, Colleen LaVinka P, Zhou F, Bergles D, Hanani M (2016) Coupled Activation of Primary Sensory Neurons Contributes to Chronic Pain. Neuron 91:1–12.

Kurtz R, Fricke M, Kalb J, Tinnefeld P, Sauer M (2006) Application of multiline two-photon microscopy to functional in vivo imaging. J Neurosci Methods 151:276–286.

Lillis KP, Eng A, White JA, Mertz J (2008) Two-photon imaging of spatially extended neuronal network dynamics with high temporal resolution. J Neurosci Methods 172:178–184.

Malmberg AB, Yaksh TL (1992) Antinociceptive Actions of Spinal Nonsteroidal Anti-Inflammatory Agents on the Formalin Test in the Rat1. 263.

Malmberg AB, Yaksh TL (1994) Voltage-sensitive calcium channels in spinal nociceptive processing: blockade of N- and P-type channels inhibits formalin-induced nociception. J Neurosci 14:4882–4890.

McCall WD, Tanner KD, Levine JD (1996) Formalin induces biphasic activity in C-fibers in the rat. Neurosci Lett 208:45–48.

Murray CW, Cowan A, Larson AA (1991) Neurokinin and NMDA antagonists (but not a kainic acid antagonist) are antinociceptive in the mouse formalin model. Pain 44:179–185.

Ottoni EB (2000) EthoLog 2.2: a tool for the transcription and timing of behavior observation sessions. Behav Res Methods Instrum Comput 32:446–449.

Pirec V, Laurito CE, Lu Y, Yeomans DC (2001) The Combined Effects of N-type Calcium Channel Blockers and Morphine on A Versus C Fiber Mediated Nociception. Anesth Analg:239–243.

Puig S, Sorkin LS (1996) Formalin-evoked activity in identified primary afferent fibers: systemic lidocaine suppresses phase-2 activity. Pain 64:345–355.

Rausell E, Cusick CG, Taub E, Jones EG (1992) Chronic deafferentation in monkeys differentially affects nociceptive and nonnociceptive pathways distinguished by specific calcium-binding proteins and down-regulates gamma-aminobutyric acid type A receptors at thalamic levels. Proc Natl Acad Sci U S A 89:2571–2575.

Reddy GD, Saggau P (2005) Fast three-dimensional laser scanning scheme using acousto-optic deflectors. J Biomed Opt 10:64038.

Sekiguchi KJ, Shekhtmeyster P, Merten K, Arena A, Cook D, Hoffman E, Ngo A, Nimmerjahn A (2016) Imaging large-scale cellular activity in spinal cord of freely behaving mice. Nat Commun 7.

Shoham S, O’Connor DH, Segev R (2006) How silent is the brain: is there a “dark matter” problem in neuroscience? J Comp Physiol A Neuroethol Sens Neural Behav Physiol 192:777–784.

Simons SB, Escobedo Y, Yasuda R, Dudek SM (2009) Regional differences in hippocampal calcium handling provide a cellular mechanism for limiting plasticity. Proc Natl Acad Sci 106:14080–14084.

Smith-Edwards KM, DeBerry JJ, Saloman JL, Davis BM, Woodbury CJ (2016) Profound alteration in cutaneous primary afferent activity produced by inflammatory mediators. Elife 5.

Stosiek C, Garaschuk O, Holthoff K, Konnerth A (2003) In vivo two-photon calcium imaging of neuronal networks. Proc Natl Acad Sci U S A 100:7319–7324.

Sun XR, Badura A, Pacheco DA, Lynch LA, Schneider ER, Taylor MP, Hogue IB, Enquist LW, Murthy M, Wang SS-H (2013) Fast GCaMPs for improved tracking of neuronal activity. Nat Commun 4:2170.

Thériault G, Cottet M, Castonguay A, McCarthy N, De Koninck Y (2014) Extended two-photon microscopy in live samples with Bessel beams: steadier focus, faster volume scans, and simpler stereoscopic imaging. Front Cell Neurosci 8:139.

Tian L, Hires SA, Mao T, Huber D, Chiappe ME, Chalasani SH, Petreanu L, Akerboom J, McKinney SA, Schreiter ER, Bargmann CI, Jayaraman V, Svoboda K, Looger LL (2009) Imaging neural activity in worms, flies and mice with improved GCaMP calcium indicators. Nat Methods 6:875–881.

Usoskin D, Furlan A, Islam S, Abdo H, Lönnerberg P, Lou D, Hjerling-Leffler J, Haeggström J, Kharchenko O, Kharchenko P V, Linnarsson S, Ernfors P (2014) Unbiased classification of sensory neuron types by large-scale single-cell RNA sequencing. Nat Neurosci 18.

Vaccarino AL, Melzack R (1989) Analgesia produced by injection of lidocaine into the anterior cingulum bundle of the rat. Pain 39:213–219.

Westenbroek RE, Hoskins L, Catterall WA (1998) Localization of Ca2+ channel subtypes on rat spinal motor neurons, interneurons, and nerve terminals. J Neurosci 18:6319–6330.

Williams EK, Chang RB, Strochlic DE, Umans BD, Lowell BB, Liberles SD (2016) Sensory Neurons that Detect Stretch and Nutrients in the Digestive System. Cell 166:209–221.

Wu A, Dvoryanchikov G, Pereira E, Chaudhari N, Roper SD (2015) Breadth of tuning in taste afferent neurons varies with stimulus strength. Nat Commun 6:8171.

Yamamoto T, Yaksh TL (1992) Comparison of the antinociceptive effects of pre- and posttreatment with intrathecal morphine and MK801, an NMDA antagonist, on the formalin test in the rat. Anesthesiology 77:757–763.

Zariwala HA, Borghuis BG, Hoogland TM, Madisen L, Tian L, De Zeeuw CI, Zeng H, Looger LL, Svoboda K, Chen T-W (2012) A Cre-dependent GCaMP3 reporter mouse for neuronal imaging in vivo. J Neurosci 32:3131–3141.

